# Macrophage phenotype transitions in a stochastic gene-regulatory network model

**DOI:** 10.1101/2022.10.21.513139

**Authors:** Anna-Simone Frank, Kamila Larripa, Hwayeon Ryu, Susanna Röblitz

**Affiliations:** University of Bergen, Computational Biology Unit, Department of Informatics, Bergen, Norway; California State Polytechnic University Humboldt, Department of Mathematics, Arcata, California, U.S.A.; Elon University, Department of Mathematics and Statistics, Elon, North Carolina, U.S.A.

## Abstract

Polarization is the process by which a macrophage cell commits to a phenotype based on external signal stimulation. To know how this process is affected by random fluctuations and events within a cell is of utmost importance to better understand the underlying dynamics and predict possible phenotype transitions. For this purpose, we develop a stochastic modeling approach for the macrophage polarization process. We classify phenotype states using the Robust Perron Cluster Analysis and quantify transition pathways and probabilities by applying Transition Path Theory. Depending on the model parameters, we identify four bistable and one tristable phenotype configuration. We find that bistable transitions are fast but their states less robust. In contrast, phenotype transitions in the tristable situation have a comparatively long time duration, which reflects the robustness of the states. The results indicate parallels in the overall transition behavior of macrophage cells with other heterogeneous and plastic cell types, such as cancer cells. Our approach allows for a probabilistic interpretation of macrophage phenotype transitions and biological inference on phenotype robustness. In general, the methodology can easily be adapted to other systems where random state switches are known to occur.

## Introduction

Macrophages are a specific type of immune cells^1^. In a process called polarization macrophages adopt different phenotypes in response to (external) signals and their microenvironment. Such stimulation results in a continuum of macrophage phenotype variety and functionality that has been simplified into a dichotomous framework: On one end, there are classically activated (M1-phenotype) macrophages and on the other end alternatively activated types (M2-phenotype). M1-like macrophages are pro-inflammatory and direct host defense, whereas M2-like macrophages have quite the opposite effect– they resolve inflammation and repair tissue^2^. In between these extremes, unique or mixed phenotypes have been observed^3^, particularly in pathological contexts such as neurodegenerative disorders^4^ or the tumor microenvironment^5^.

Macrophage polarization is regulated by cytokine signals which induce the Jak-STAT pathway in these cells^6^. This pathway communicates information from chemical signals outside of a cell to the nucleus, resulting in the activation of genes. As an example, STAT1 activation facilitates the transcription of genes which create pro-inflammatory products^7^. Within the binary phenotype classification system, simultaneous high STAT1 and low STAT6 activation is interpreted as the M1 phenotype, whereas reversed activation levels characterize the M2 phenotype^8^.

We furthermore know that macrophages within a tumor can switch between phenotypes, which determine their functionality to be either promoting or suppressing to tumor growth^9^. In other words, macrophages are very plastic cells^10^ that can reverse their activation state fully and rapidly. Therefore, a given macrophage may participate sequentially in the induction and resolution of inflammation^11^.

Even with this biological insight, knowledge gaps exist in understanding how microenvironmental signaling drives polarization, phenotype switching and functionality, and ultimately pathological outcomes.

To address these knowledge gaps, several deterministic model frameworks, based on Ordinary Differential Equations (ODEs), have been developed in recent years to better understand macrophage polarization processes, and the transitions between phenotypes, e.g.,^12–15^. These models confirmed the existence of already known phenotypes, and predicted others not yet described in the literature. Furthermore, bifurcation analysis revealed the possibility of phenotype switches that are known to exist in living cells and cell cultures^3^.

Deterministic model frameworks build on the assumption that there are a large number of molecule (e.g., transcription factors) present within a cell. Contrary to this assumption, (human) cells can include only a restricted number of molecule copies, such that intrinsic noise and random fluctuations dominate.

A deeper knowledge and better understanding of the macrophage polarization process within the stochastic cell environment is therefore necessary for the development of new therapeutic approaches which maintain or reverse the macrophage phenotype functionality. However, such innovations are conditioned on our knowledge of the possible pathways and respective probabilities of phenotype switches, as well as the responsible underlying mechanisms. Such knowledge is currently missing.

In this paper, we therefore aim to mimic a stochastic cell environment by taking into account randomness within a single macrophage cell. With this approach, we first study possible phenotype configurations and compare them to the outcome of deterministic model approaches, to then analyze the transition routes of these phenotypes and their corresponding probabilities.

The article is organized in the following way. We start with the presentation of the methodological approach, followed by the results, and end with their discussion. The Supplementary Material online includes more details on the methods, as well as additional figures and tables describing our results. Implementation codes (in Matlab) of the methodology are available. A large contribution of this work is that the methodologlical approach can be applied to multi-stable systems beyond macrophages.

## Methods

An overview of the methodological approach used in this article is presented in Fig. 1.

**Figure 1.**
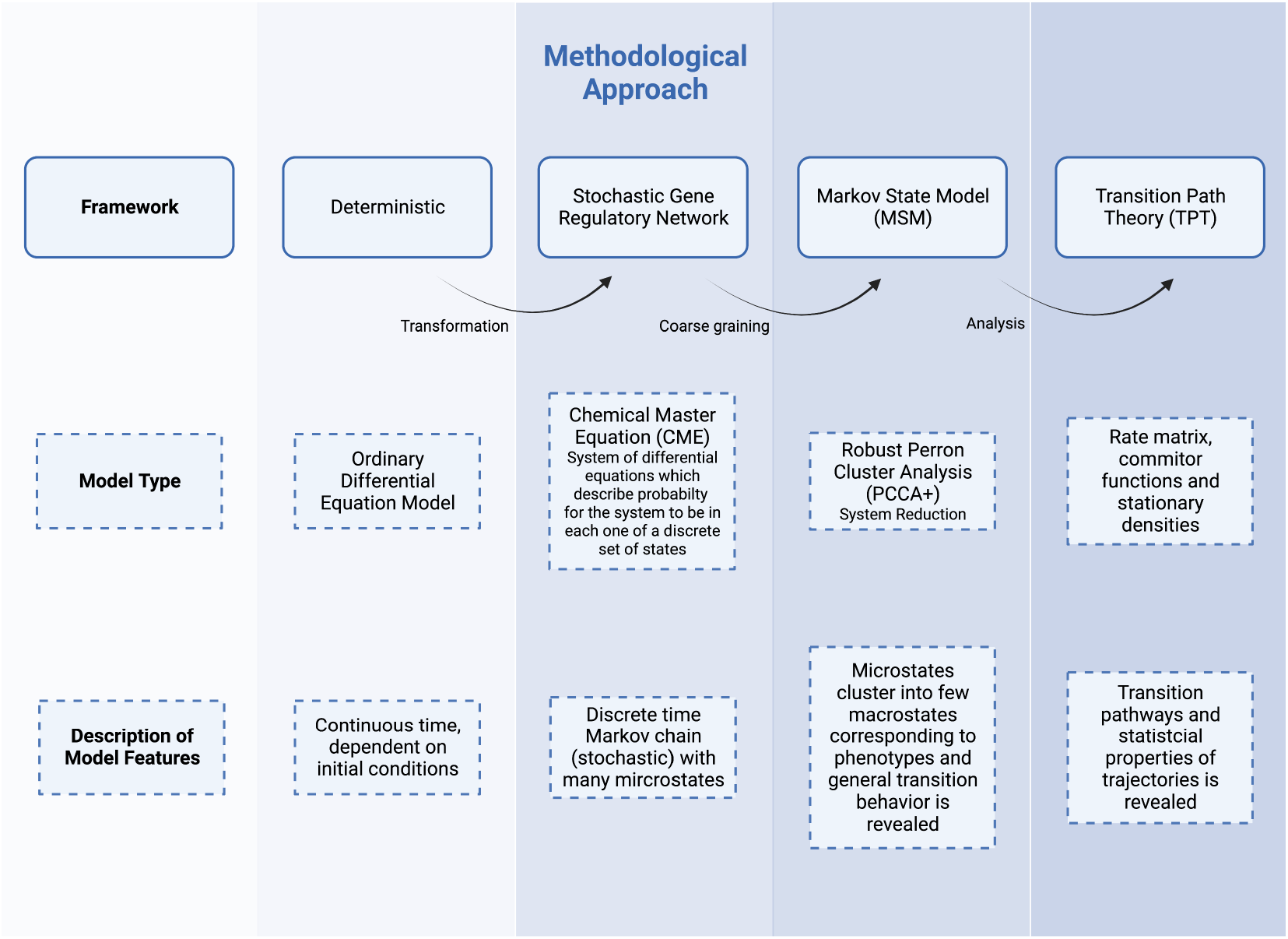
Flowchart of the methodological approach. First, a deterministic model of macrophage polarization is transformed into a stochastic modeling framework. We then use a clustering approach to identify possible multi-stable phenotypes based on several different parameter sets. Finally, to calculate phenotype transition paths and probabilities between the identified phenotypes, we apply Transition Path Theory. Created with Biorender.com.

### Building the stochastic gene-regulatory network model

The Chemical Master equation (CME) is a fundamental description of interacting molecules commonly used to model noisy gene regulatory networks. It is a system of differential equations that describes the evolution of *probability densities* for Markov processes when a system jumps from one discrete state to another in continuous time.

Let *p*(*i, j*) denote the probability to find the system in state (*x*_1_ = *i, x*_2_ = *j*) at time *t*, whereby *x*_1_ represents STAT1 and *x*_2_ represents STAT6, and *i, j* ∈ ℕ are their respective copy numbers. Then, based on the two-dimensional deterministic ODE model in^15^, which is sketched in Fig. 2, we define our stochastic model (i.e., the CME) as

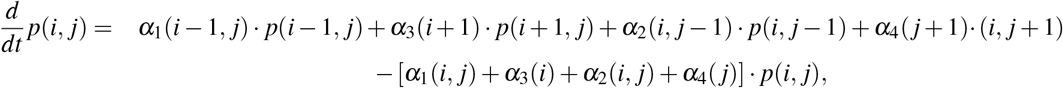

with propensity functions

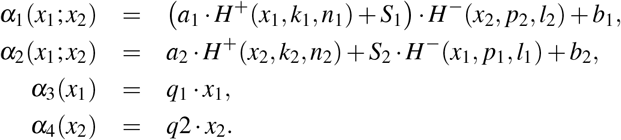

**Figure 2.**
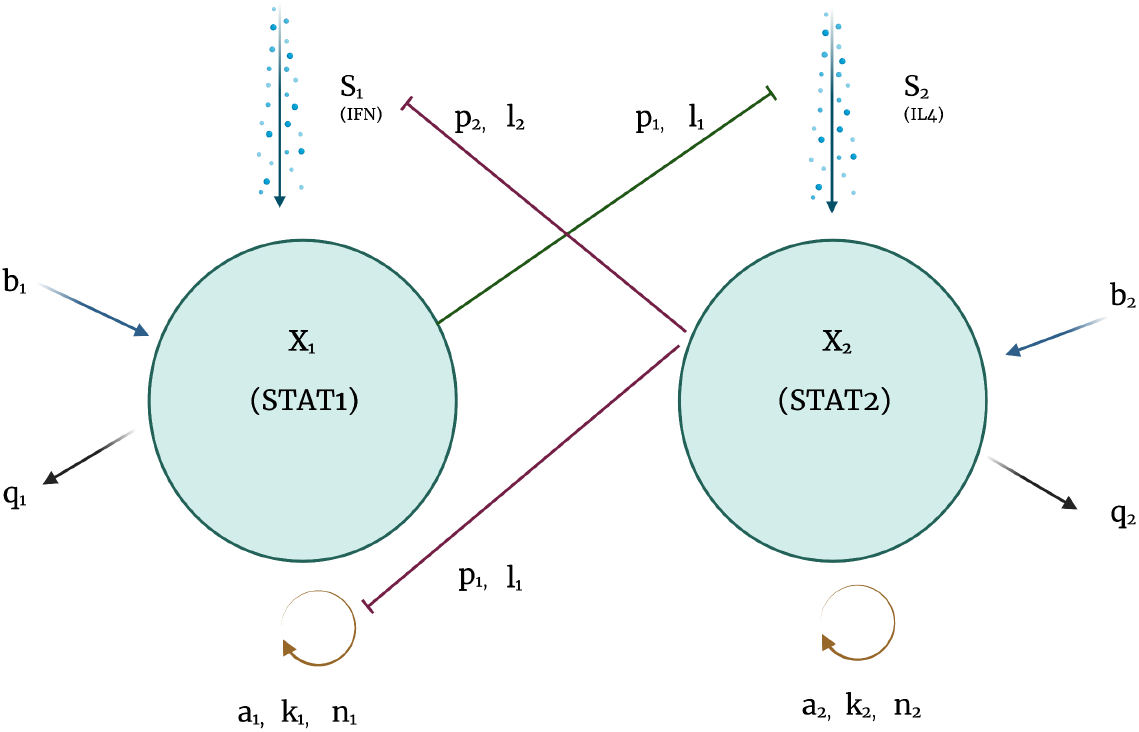
Schematic diagram of the modeled signaling network. The variable *x*_1_ represents STAT1, a marker for M1 polarization, and the variable *x*_2_ represents STAT6, a marker for M2 polarization, along with two input signals (denoted by *S*_1_ and *S*_2_). Self-stimulation of STATs (*a_i_, n_i_, k_i_, i* ∈ {1, 2}) is represented by the orange loops at the bottom of the figure, while processes of mutual-inhibition are depicted with red and green inhibiting lines (*p_i_, l_i_, i* ∈ {1, 2}). The incoming blue arrows (*b_i_, i* ∈ {1, 2}) represent basal activation of STATs which also occurs in the absence of cytokine signalling. The outgoing black arrows (*q_i_, i* ∈ {1, 2}) represent deactivation of STATs. Incoming arrows at the top represent cytokine signalling (IFN (*S*_1_) and IL4 (*S*_2_) respectively). Note the asymmetry in that STAT6 inhibits both the input signal and self stimulation of STAT1, but STAT1 impacts only the input signal for STAT6. See^15^ for a detailed description of the model equations. Created with Biorender.com.

The term 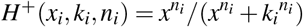 denotes a stimulatory Hill function, whereas 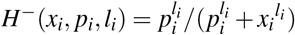 denotes an inhibitory Hill function, see^15^ for a biological rationale for their use.

If the state space is enumerated with states *k* = 1, 2,…, *n*, this equation can be written in matrix notation,

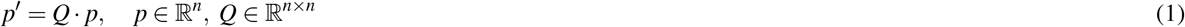

whereby the transition rate matrix *Q* has column sums zero. The stationary density *π* ∈ ℝ^*n*^ is given by the eigenvector of *Q* corresponding to eigenvalue *λ* = 0,

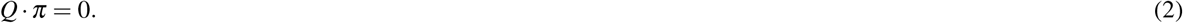

### Phenotype identification via clustering

Stochastic changes in particular patterns of gene expression have been identified with spontaneous phenotype transitions that can diversify otherwise identical cell-populations. These gene expression patterns correspond to metastable regions in the state space of the CME. The dynamics remains in these regions for a long time before it rapidly switches to another metastable region.

Multistability is characterized by the occurrence of a cluster of *n_c_* > 1 eigenvalues 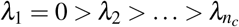 close to zero for the matrix *Q*. Note that the eigenvalue *λ*_1_ = 0 is equivalent to the Perron root *λ*_1_ = 1 (also called the leading eigenvalue or dominant eigenvalue) of the corresponding transition probability matrix *P*(*τ*) = exp(*τ·Q*). It has been demonstrated previously^16^ that the subspace spanned by the corresponding eigenvectors, *X*, contains all information about the location of the metastable regions. To extract this information, spectral clustering approaches, such as the Robust Perron Cluster Analysis (PCCA+)^16^, are useful. The PCCA+ computes a transformation matrix *A* such that the transformed eigenvectors *χ* = *X·A* become so-called membership vectors. These membership vectors form a fuzzy clustering of the state space in that they assign to any state the probabilities for belonging to any of the *n_c_* clusters, which represent the metastable regions. In the context of macrophage polarization, the metastable regions define the different phenotypes.

Furthermore, the membership vectors are used to project (coarse grain) the dynamics onto the space of metastable regions by

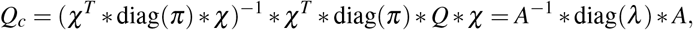

where 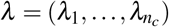.

While the coarse grained transition rates might be difficult to interpret, the corresponding coarse grained transition probabilities for a time interval of length *τ* can be obtained via the matrix exponential

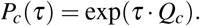

The statistical weights *w_k_* of the clusters can be computed by

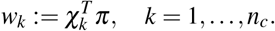

Similarly, the normalized partial densities of the clusters are given by

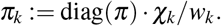

### Explorations of parameter sets

While for the ODE model^15^ three different parameter sets gave rise to bi-, tri-, and quad-stability, the parameter sets leading to multistability in the stochastic environment might not be necessarily the same.

We therefore used several approaches to determine the parameter sets yielding multistability in the stochastic setting:

First, we transformed the parameters from the ODE model^15^ from concentration units into copy numbers, as described in the Supplementary Methods online, and used the derived copy numbers as parameter values for the stochastic model.

Second, the most sensitive parameters in the ODE model^15^ were varied and their variations served as parametrization for the stochastic model.

Third, we performed a Latin Hypercube Sampling (LHS) approach to further explore the parameter space of the stochastic model (details described in the Supplementary Methods online).

Fourth, we manually sampled the parameter values based on the derived copy numbers from approach one.

Besides the criterium for multistability (i.e., a spectral gap in the *zero* eigenvalue spectrum of the transition rate matrix *Q*), the resulting parameter sets were also assessed based on the quality of the clustering (the crispness of the membership functions *χ*) and a visual (spatial) separation of the resulting metastable regions.

### Computing phenotype transition paths and probabilities via Transition Path Theory

Transition Path Theory (TPT) has originally been developed in the context of molecular dynamics simulations, but has previously also been applied to gene regulatory networks^17, 18^. The theory is described in detail in numerous papers^19–21^.

Based on the identified macrophage phenotypes, TPT is applied to determine the transition pathways a long which macrophages switch their phenotype, together with the corresponding path probabilities. For completeness, we briefly summarize the TPT concepts that are important for our application to macrophage polarization, but refer the reader to the Supplementary Material online, as well as the above mentioned papers for more details.

TPT takes as input the transition rate matrix 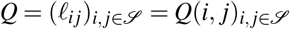 (from equation (1)) of a Markov jump process {*X*}_*t*∈ℝ_ defined on a discrete state space 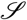. Based on the assumption that the process {*X*}_*t*∈ℝ_ is irreducible and ergodic, the time-reversed process 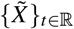 is also a Markov jump process with a rate matrix 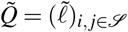 given by 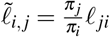^19^, where *π* is the invariant distribution (or stationary density) of both processes {*X*}_*t*∈ℝ_ and 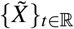, satisfying equation (2).

TPT is based on *reactive trajectories*, which are defined as transitions between two non-empty, disjoint subsets *A* and *B* of the state-space 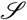, where one set is the source and the other one is the sink state^19, 20^.

As a first step, we compute the discrete forward *q*^+^ and backward *q*^−^ committor functions, as they are essential to calculate statistical properties and dynamical information of the reactive trajectories. *q*^+^ is the probability that the process that starts in 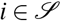 will first reach subset *B* rather than *A*, while *q*^−^ describes the probability that the process first visited sub-set *A* rather than *B*^19^. In a second step, we calculate statistical properties that allow for a dynamical interpretation of the ensemble of reactive trajectories between subsets *A* and *B* in the state space 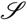 ^22^. Supplementary Table S1 online summarizes important measures in TPT to describe dynamical information and states the corresponding interpretation. For the detailed computation of these quantities, the reader is referred to the Supplementary Methods online.

In a third step, we coarse-grain the probability current of reactive trajectories, 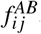, by projecting it onto the metastable state regions *S_i_* and *S_j_* from the Markov State Model, i.e., onto the phenotypes,

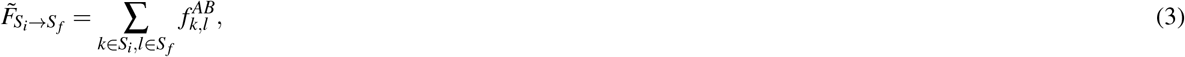

which in its normalized form reads

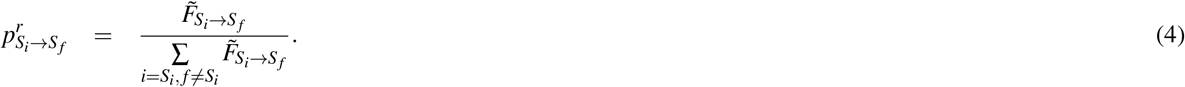

In this step, we define a set *S_k_* as the support of a partial density *π_k_* in such a way that it carries at least (1 − *ε*)·100% of this density, i.e.

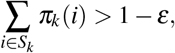

whereby we choose *ε* = 10^−6^.

Finally, by using probability tree diagrams, the relative probabilities of the path segments 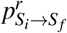 are combined along an individual pathway *P_i_* from *A* to *B* to define the *total relative probability* 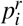 *of path P_i_*.

## Results

### Parameter sets for multistability

From the above described parameter explorations, procedure one did not result in multi-stable states, but procedures two to four showed bi- and tri-stability. From these, only procedure four lead to meaningful clusters with clearly separated phenotypes. Thus, we based the subsequent analysis on the parameter sets derived from the manual sampling procedure, which lead to five different parameter sets (cases). These are presented in Table 1.

**Table 1.** Parameter sets. The table presents the five different cases of parameter sets from the manual sampling approach, which give rise to multi-stable macrophage phenotypes in the stochastic model. The values are based the model parameters in^15^. Due to the conversion to molecule numbers, parameters have been multiplied with the factor *u·V·n_A_* = 12.9 (see Supplementary Methods online for more details on the conversion).

| Parameters | $a_1$ | $a_2$ | $b_1$ | $b_2$ | $n_1$ | $n_2$ | $k_1$ | $k_2$ | $l_1$ | $l_2$ | $p_1$ | $p_2$ | $q_1$ | $q_2$ | $S_1$ | $S_2$ |
| --- | --- | --- | --- | --- | --- | --- | --- | --- | --- | --- | --- | --- | --- | --- | --- | --- |
| Case 1 (bistable) | 5 | 5 | 15 | 0.05 | 6 | 6 | 1 | 1 | 1 | 1 | 0.5 | 1 | 5 | 1 | 2 | 2/12.9 |
| Case 2 (bistable) | 15 | 5 | 0.05 | 0.05 | 6 | 6 | 1 | 1 | 1 | 1 | 0.5 | 1 | 5 | 1 | 2 | 2/12.9 |
| Case 3 (bistable) | 15 | 5 | 0.05 | 0.05 | 6 | 6 | 1 | 1 | 1 | 1 | 0.5 | 1 | 5 | 5 | 1 | 1/12.9 |
| Case 4 (bistable) | 5 | 5 | 0.001 | 0.05 | 6 | 6 | 1 | 1 | 1 | 1 | 0.5 | 1 | 5 | 1 | 2 | 2/12.9 |
| Case 5 (tristable) | 15 | 5 | 0.05 | 0.05 | 6 | 6 | 1 | 1 | 1 | 1 | 0.5 | 1 | 5 | 1 | 4 | 4/12.9 |

Parameter sets leading to bi- and tristability are notably different from each other, see Table 1. For example, the phenotype observation in case 1 is driven by the parameter ratio of *b*_1_ ≫ *b*_2_, and partly by *q*_1_ > *q*_2_. Tristability (case 5) results from bistability (case 2) by increasing the external input signals *S*_1,2_. Other bistable cases (e.g., case 3 and 4) emerge, respectively, from parameter cases 2 and 1 by a change of values in *a*_1_, *b*_1_, *q*_2_ and *S*_1,2_. These parameter dependencies potentially reflect the differences in cellular conditions driving phenotype emergence.

### Identified phenotypes in the stochastic environment

We observe diversity in multistable macrophage phenotype configurations, specifically bi- and tristability, as presented by the clustering results in Supplementary Table S3 online. There, multistability is also visible in the corresponding transition probability matrices *P_c_* and the surface plots of the partial densities, which represent the multiple phenotypes states (see Fig. 3 for bistability and Supplementary Fig. S1 online for tristability).

**Figure 3.**
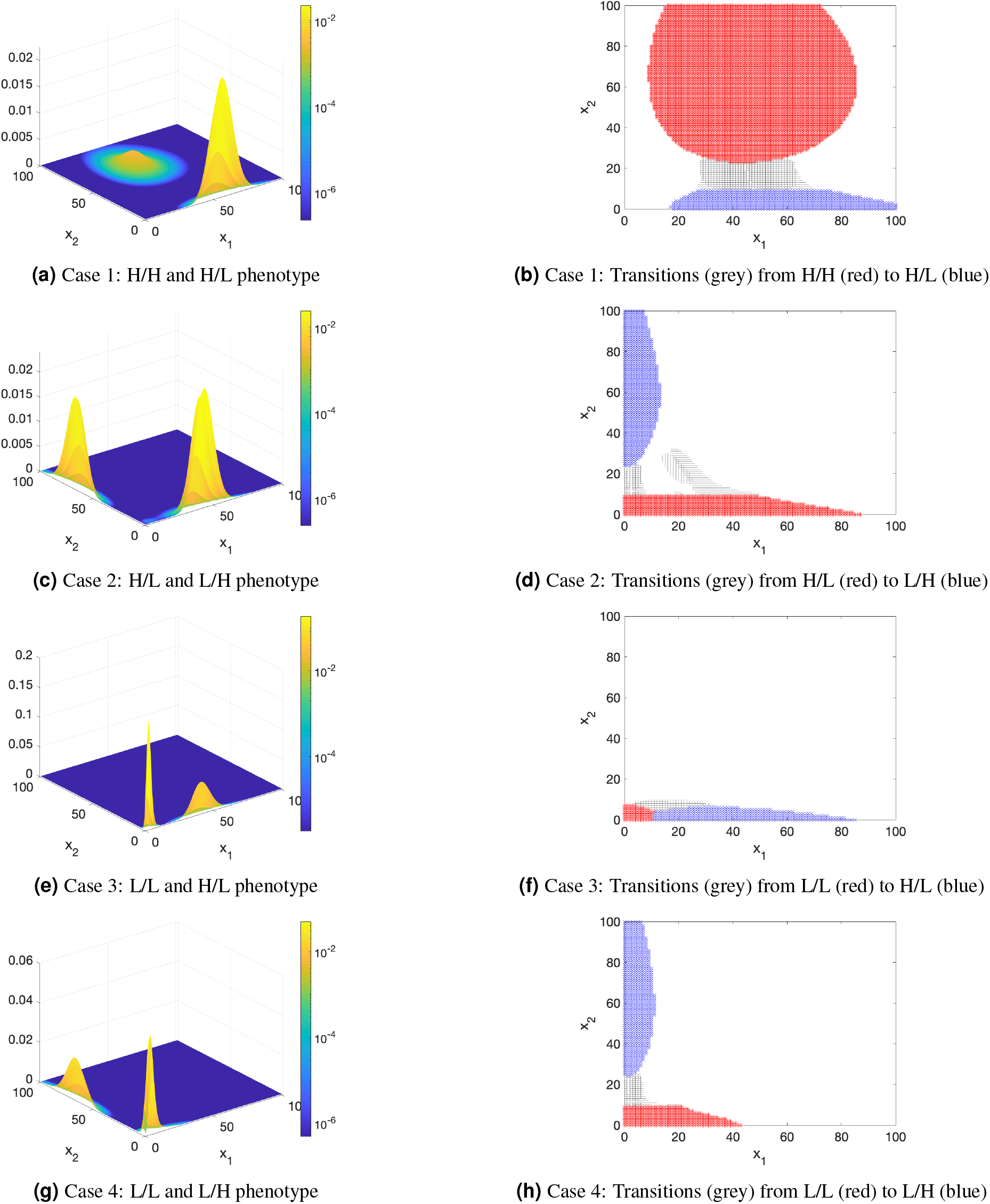
Bistable phenotype configurations. The left panel shows a surface plot of the sum of the partial densities which correspond to two metastable clusters. The ratio between *x*_1_, *x*_2_ defines the phenotypes. In (a) we see high activation levels in both *x*_1_ and *x*_2_, and thus can classify a High/High (H/H) phenotype. High activation in *x*_1_ and low activation in *x*_2_, describes a H/L phenotype. Similarly, we see bistable clusters of phenotypes in (c) H/L and L/H, in (e) L/L and H/L and in (g) L/H and L/L. The right panel shows calculated transition path flow directions in (b), (d), (f) and (h). In these transition flow graphs, the red colored set indicates the starting set and the blue colored set the ending set. The gray colored area represents the transition flow. The Supplementary Table S4 online gives a quantitative overview over the transition dynamics, including the respective paths and probabilities.

Depending on parameters used in the model (see Table 1), Fig. 3 shows four different pairs of bi-stable phenotype sets, which are defined based on the relative levels (e.g., low, high) of activation in *x*_1_/*x*_2_: (1) high *x*_1_/low *x*_2_ (H/L) in Fig. 3a and high *x*_1_/high *x*_2_ (H/H), (2) low *x*_1_/high *x*_2_ (L/H) and high *x*_1_/low *x*_2_ (H/L) in Fig. 3c, (3) low *x*_1_/low *x*_2_ (L/L) and high *x*_1_/low *x*_2_ (H/L) in Fig. 3e and (4) low *x*_1_/low *x*_2_ (L/L) and low *x*_1_/high *x*_2_ (L/H) in Fig. 3g.

Only one case with tri-stable phenotypes was detected, which comprised the L/L, the H/L and the L/H phenotypes (see Supplementary Fig. S1 online).

Furthermore, the statistical weights *w_i_* (*i* = 1, 2, 3), in Supplementary Table S3 online are an indication for how much time the dynamical process spends in a phenotype state before switching to another one. The higher the weight, the longer the expected holding time.

### Phenotype transitions in the bistable case

The dynamic flow paths between start (red colored) and end (blue colored) sets, where each set represents a specific phenotype, are presented in Fig. 3 and Supplementary Fig. S2 online.

What the information above and in Supplementary Table S4 online reveal is that for the parameter set in case 4 (see Table 1), a macrophage will never transition directly from L/L to L/H or vice-versa (0% probability) but will always pass through the complement set (C) when making this phenotype change. However, in Case 3, we see that a macrophage is much more likely to directly transition between L/L and H/L rather than meander through a complement set by considering the transition probabilities. Additionally, the transition time can be compared between different cases (see Supplementary Table S4 online). For example, it is an order of magnitude faster for a macrophage to transition from L/L to H/L in Case 3 rather than from H/L to L/L.

Overall, the transitions flow mostly (nearly 100%) on one specific path, directly from the start to the end set, through the complement set (C). This observation is somewhat different from what we observe when there are more than two phenotypes present (see the results on the tristable case below).

### Phenotype transitions and probabilities in the tristable case

The dynamic flow paths between the three identified phenotypes (L/L, H/L and L/H) are presented in Fig. 4 and Supplementary Fig. S3 online for different combinations of start (red color) and end sets (blue color).

**Figure 4.**
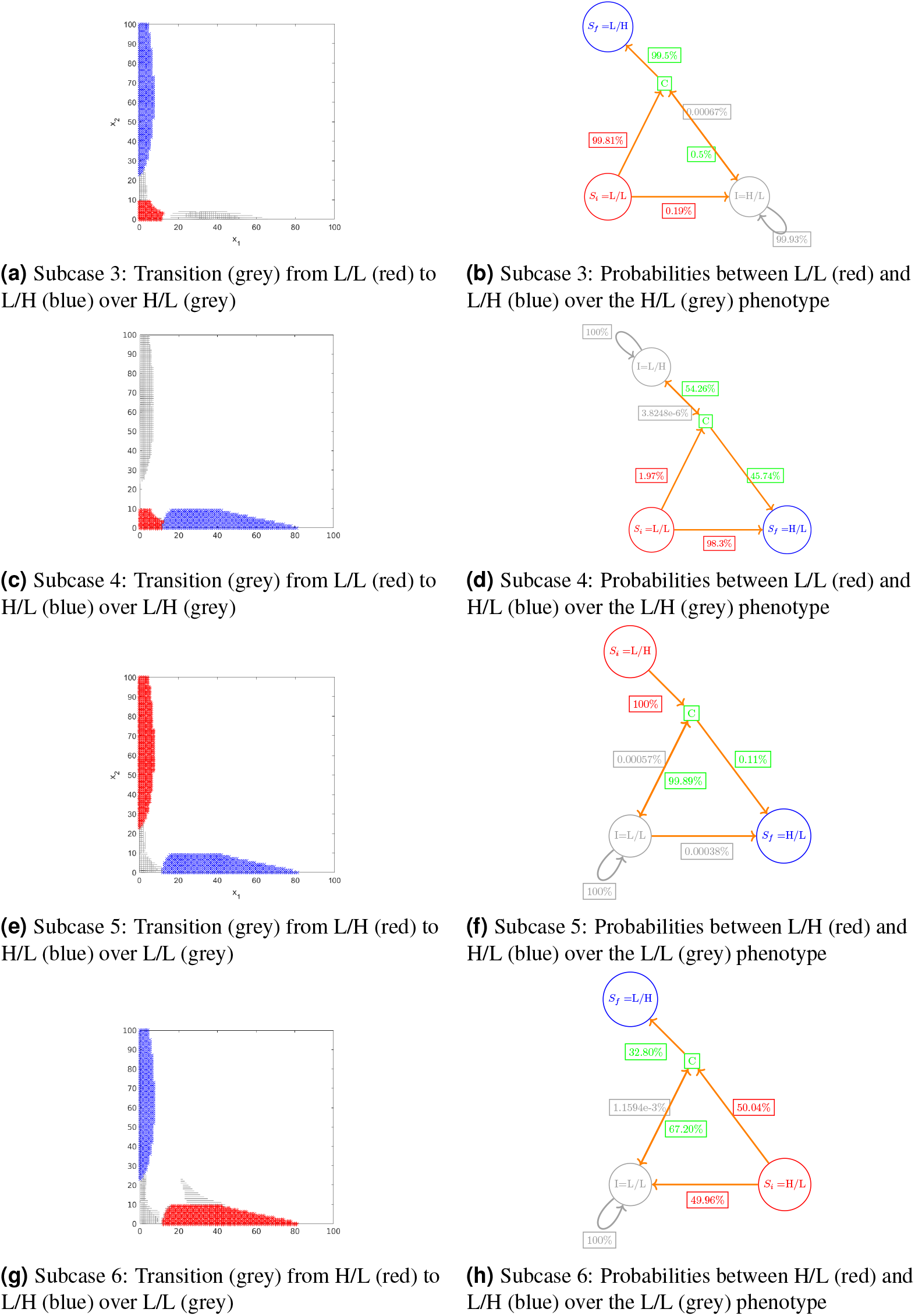
Transition flows, paths and probabilities between three phenotypes. The left column shows transition path flow (in grey color) in the CME state-space between a given start set (in red color) and end set (in blue color). The right column presents the transition network together with the corresponding path segment probabilities, added next to the arrows of the flow direction. Red colored circles represent the start sets (*S_i_*, with i for initial), and blue ones the end sets (*S_f_*, with f for final). Intermittent phenotype states (I) are colored in grey and green represents the state space without any phenotype, i.e., the complement set (C) to the states (*S_i_, IandS_f_*). The path probabilities were calculated from equation (4) in the Methods section. Their color refers to the set from where the transition flow comes. Supplementary Table S5 online gives a quantitative overview over the transition dynamics, including all possible transition paths and their respective probabilities. Subcase 1 and 2 with combined end and start states are presented in Supplementary Fig. S3 online.

The transition networks are illustrated in Fig. 4b, 4d, 4f, and 4h as well as in Supplementary Fig. S3.b and S3.d online. As it can be seen from these figures, the choice of start and end set has an important role for the transition dynamics and phenotype realizations of the macrophage cell. This choice influences the specific transition paths and their probabilities. In the following, we therefore distinguish between the different flow directions by enumerating them as ‘subcases 1–6’.

In subcase 1, the states L/H and H/L are the end sets and L/L is the start set (see Supplementary Fig. S3.a and S3.b online), while in subcase 2 the start sets is defined as the combination of the L/L and H/L phenotype (see Supplementary Fig. S3.c and S3.d online). In these two cases, we observe short transition times (*t* = 0.16 and 0.40, respectively) (see Supplementary Table S5 online) with high transition probabilities for the different paths. Particularly, subcase 1 shows that cells are more likely to end up in the H/L phenotype (57.4%) than in the L/H phenotype (43.6%). In subcase 2, 90.3% of the transitions to the L/H state start in the L/L phenotype, while only 9.6% start in the H/L state.

Both subcases 3 and 4 start from the same start set (L/L), but their end phenotype state as well as the intermittent phenotype state differ (see Fig. 4a and 4c). For example, in subcase 4 macrophage cells coming from the L/L state end up with 99.2% probability in the H/L phenotype (either directly or via the complement, see Supplementary Table S5 online). There is, however, an additional very small probability that the macrophage switches to the L/H phenotype, and stays there for a very long time, before settling in the final H/L state. The stay in the L/H state is represented by the recursive (gray) flow arrow in the corresponding transition network (see Fig. 4d). This discourse over the L/H phenotype, with a very low segment probability (3.8e-6%) of leaving the state (L/H), makes the transition from the L/L to the H/L phenotype lasting very long on average (*t* = 135337) and thus very unlikely (see Supplementary Table S5 online). Similar interpretations can be drawn for subcase 3 based on the presented results in Fig. 4a and 4b and the Supplementary Table S5 online. Here, however, the results indicate that the macrophage cell is able to leave the intermittent state (H/L). This is also indicated by the relatively fast transition time (*t* = 2.03).

Subcases 1-3 are examples in which a macrophage cell successfully switches its phenotype (i.e., the majority of the transition flow reaches the defined end phenotype within reasonable time).

Similar to subcase 4, also subcases 5 and 6 represent a situation where the final phenotype state will probably not be reached. Here, the start sets are the L/H, respectively the H/L phenotype, and the end sets are the H/L and L/H states, respectively (see Fig. 4e and Fig. 4g). The dynamic information in Supplementary Table S5 online, as well as the segment probabilities in the corresponding transition networks in Fig. 4f and Fig. 4h indicate that the transition flow is recurrently drawn towards the intermittent phenotype state (I) with high probability (i.e., there are a very low probability path segments for leaving the intermittent state, while the path segment probabilities leading to this state are high). In other words, it will require several transition iterations between the intermittent state (I) and the complement set (C) until the end state is reached. This observation can also be inferred from the low probabilities of the respective transition paths (see Supplementary Table S5 online). The average length of a successful transition (*t* = 699 and *t* = 1359, respectively, see Supplementary Table S5 online) reflects this result as well.

## Discussion

Informed knowledge about phenotype states, paths and their probabilities in a realistic setting is crucial to predict the role and functionality of macrophage cells in disease progression and to understand the underlying mechanisms better. Until now, however, it was unclear how macrophages “behave” in their stochastic environment. Therefore, we designed a methodological approach to mimic the macrophage switching dynamics in a stochastic cellular environment. We show that phenotype multistability is maintained in the stochastic setting, but that the number and combinations of emerging phenotypes differ from a deterministic approach, as compared to the results in Frank et al.^15^. This illustrates the influence and sensitivity of single-cell decisions to random fluctuations, which is in line with prior work by^17,18,23^, where the authors analyzed how stochasticity affects cell-fate-decisions of cancer, embryonic stem cells or closed gene regulatory network motifs. In addition, we gained first insights into the probabilistic nature of macrophage phenotype switches, which depends on the number of phenotypes, the specific type, as well as the switch direction.

In particular, our analysis revealed that for different parameter configurations, different phenotypes (e.g., L/L, H/L, L/H and H/H) emerge. These phenotypes coincide with those identified in^15^. However, our results also indicate that the number of multi-stabile phenotypes (e.g., quad-stable) in stochastic systems is restricted to bi- and at most tri-stability, which is in accordance with the current literature^23–25^. The validity of this hypothesis would have a huge impact on our understanding of macrophage cells and needs to be confirmed. Furthermore, the biological mechanisms behind changes in different phenotype configurations need more attention, especially as macrophage polarization is dynamic across time^26^. For example, our observation that specific parametric sets leading to different phenotype outcomes can evolve from one another or in some cases remain mutually exclusive should be replicated in laboratory experiments, and their kinetic parameters within the cells should be quantified. Such an endeavor is a crucial first step to actually reconcile the biologically observed macrophage phenotypes and computational results^18,27^. The overall lack of data on macrophage markers restricts our knowledge about corresponding kinetic parameters and consequently on macrophage subtypes and their transition behavior^27^. This restricts also our analysis and results to purely synthetic scenarios and warrants the need for more data on macrophage phenoytpe subtypes, their markers and kinetic parameters to gain better understanding of the underlying processes of these heterogeneous cells through modeling in the future.

Considering the level of activation of STAT1 and STAT6, we could interpret the L/H phenotype as M2-like, and H/L phenotype as M1-like, while H/H would represent a mixed phenotype and L/L an M0 phenotype^15^. While the M0, M1 and M2 phenotypes are known to exist, mixed phenotypes (i.e., H/H) are less common. Indeed, unique or mixed phenotypes have been observed^3^, particularly in pathological contexts such as neurodegenerative disorders^4^ or the tumor microenvironment^5^. For example, mixed phenotype macrophages have been recently observed in early-stage lung cancer^28^ as well as human breast cancer^29^ and there are open questions both about the biological mechanisms which lead to their existence and their importance in tumor progression and control^27^. Recent biological experiments in phenotype profiling^28^ have however demonstrated that tumor associated macrophages (TAMs) in early lung cancer often demonstrate a mixed phenotype, coexpressing M1 and M2 markers. This result indicates that in early-stage tumors, the phenotype of macrophages is mixed and not skewed towards either M1 or M2. It is well known that in advanced tumors, M2 macrophages are abundant^30^, indicating that a large population of macrophages undergo a phenotype switch (or switches) as the tumor progresses. Similar observations have been made in^31^. Here the authors describe a “switch phenotype” or “M3-phenotype”, which actually mixes attributes from both the M1 and M2 phenotype and has been observed in experiments when the phenotype spectrum shifted from M1 to M2 or vice versa. In our results, we have observed such a shift between the M1 to the M2 phenotypes over the complement set, which implies that the phenoytpe has to mix M1 and M2 attributes temporarily to reach the other “side”. The hypotheses made by^31^, however, indicate that another path exists that leads through a stable fourth phenotype, similar to the H/H one observed in the bistable situation, linking M1 and M2. Although our results did not confirm their claim, we cannot rule out the existence of such a phenotype and transition path. Our results also show that bistable phenotype switches are faster than switches between three phenotypes. This results coincides with the finding in^27^ that macrophage phenotypes have different half-life spans (e.g., M1 and mixed phenotypes have shorter spans), which consequently could impact the speed of specific transitions. If, for example, a transition paths passes through an intermediate phenotype that has a long half-life, the switch to the end phenotype would take longer time. In our results, such an observation was specifically clear in the M1-M2 phenotype transition over the M0 state, which lead to very long average transition duration. This duration reflects the energy barrier that has to be crossed for the phenotypes to switch states^32^. For example, the longer the time, the more energy is needed for a transition and the robuster are the phenotype states^32^. A similar observation has also been confirmed in a study on cancer cells, where the authors showed that specific phenotype configurations were maintained over a long time period^23^. In addition, the transition network probabilities in^23^ for cancer sub-populations are comparable in terms of rate and magnitude to our results in the tristable case. In particular, the study also shows that some phenotypes have a very high probability to remain in a specific state once this state has been reached. Finally, the observation that cancer stem like cells arise form non-stem-like cells, is similar to our observation of the L/L (M0-like) phenotype^23^. Although cancer and macrophage cells are not identical, they share similar attributes like heterogeneity and plasticity^33^. Thus, the observed similarities, between our study and the results by Gupta et al.^23^, in transition dynamics might be another common attribute of these cell types. This inference however needs further investigation. In addition, future studies should investigate if the transition probabilities for single macrophage cells coincide with data from in-vivo and in-vitro experiments and how these probabilities are altered in populations of macrophage cells. Particularly, the presented methodological approach in this manuscript could be of use for such future work.

Overall, we have shown that our methodological approach is a useful tool to uncover the underlying dynamic differences in multi-stable systems and to test different hypotheses about potential configuration states. Upon the availability of more data on macrophages and their kinetic parameters, we believe that this approach will be of great importance to further investigate and clarify our understanding of the mechanisms behind particular cell transition pathways.

## Acknowledgements

The research of SR was funded by Trond Mohn Foundation, Grant No. BFS2017TMT01.

The authors are grateful for the support of the American Institute of Mathematics through their Structured Quartet Research Ensemble funding.

## Author contributions statement

All authors conceptualized and designed the work as well as performed the analysis. A.S.F. and S.R. implemented the software used in this work. All authors drafted the manuscript, edited and reviewed the manuscript and approve its submitted version. All authors agree that they are personally accountable for their own contributions and ensure accuracy and integrity of any parts of the work.

## Additional information

### Accession codes

The Matlab codes for all presented cases are available on GitHub (https://github.com/a-sfrank/macro-tpt.git). The cPCCA+ Matlab code, as described in^34^, was downloaded from GitHub (https://github.com/sroeblitz/cPCCA.git), adapted and integrated into our implementation.

## References

1. Lahmar, Q. et al. Tissue-resident versus monocyte-derived macrophages in the tumor microenvironment. Biochim. Biophys. Acta - Rev. Cancer. 1865, 23–34 (2016).

2. Mantovani, A., Sica, A. & Locati, M. Macrophage polarization comes of age. Immunity. 23(4), 344–346 (2005).

3. Sica, A. & Mantovani, A. Macrophage plasticity and polarization: in vivo veritas. J. Clin. Investig. 122(3), 787–795 (2012).

4. Garofalo, R. S. et al. Severe diabetes, age-dependent loss of adipose tissue, and mild growth deficiency in mice lacking akt2/pkb*β*. J. Clin. Invest. 112(2), 197–208 (2003).

5. Biswas, S. K., Sica, A. & Lewis, C. E. Plasticity of macrophage function during tumor progression: regulation by distinct molecular mechanisms. J. Immunol. 180(4), 2011–2017 (2008).

6. Hu, X., Chen, J., Wang, L. & Ivashkiv, L. B. Crosstalk among Jak-STAT, Toll-like receptor, and ITAM-dependent pathways in macrophage activation. J. Leukoc. Biol. 82(2), 237–243 (2007).

7. Ma, J. et al. Glycogen metabolism regulates macrophage-mediated acute inflammatory responses. Nat. Commun. 11, 1769; 1–16 (2020).

8. Bardi, G. T., Smith, M. A. & Hood, J. L. Melanoma exosomes promote mixed M1 and M2 macrophage polarization. Cytokine. 105, 63–72 (2018).

9. Poh, A. R. & Ernst, M. Targeting macrophages in cancer: from bench to bedside. Front. Oncol. 8, 49; 1–16 (2018).

10. Biswas, S. K. & Mantovani, A. Macrophage plasticity and interaction with lymphocyte subsets: cancer as a paradigm. Nat. Immunol. 11, 889–896 (2010).

11. Porcheray, F. et al. Macrophage activation switching: an asset for the resolution of inflammation. Clin. Exp. Immunol. 142(3), 481–489 (2005).

12. Nickaeen, N., Ghaisari, J., Heiner, M., Moein, S. & Gheisari, Y. Agent-based modeling and bifurcation analysis reveal mechanisms of macrophage polarization and phenotype pattern distribution. Sci. Rep. 9, 12764; 1–14 (2019).

13. Smith, T. D., Tse, M. J., Read, E. L. & Liu, W. F. Regulation of macrophage polarization and plasticity by complex activation signals. Integr. Biol. 8(9), 946–955 (2016).

14. Zhao, C. et al. A mechanistic integrative computational model of macrophage polarization: implications in human pathophysiology. PLoS Comput. Biol. 15(11), e1007468 (2019).

15. Frank, A. S., Larripa, K., Ryu, H., Snodgrass, R. G. & Röblitz, S. Bifurcation and sensitivity analysis reveal key drivers of multistability in a model of macrophage polarization. J. Theor. Biol. 509, 110511; 1–15 (2021).

16. Röblitz, S. & Weber, M. Fuzzy spectral clustering by PCCA+: application to Markov state models and data classification. Adv. Data. Anal. Classif. 7, 147–179 (2013).

17. Chu, B. K., Margaret, J. T., Sato, R. R. & Read, E. L. Markov State Models of gene regulatory networks. BMC Syst. Biol. 11, 14; 1–17 (2017).

18. Tse, M. J., Chu, B. K., Gallivan, C. P. & Read, E. L. Rare-event sampling of epigenetic landscapes and phenotype transitions. PLoS Comput. Biol. 14(8), e1006336 (2018).

19. Metzner, P., Schütte, C. & Vanden-Eijnden, E. Transition path theory for Markov jump processes. Multiscale Model. Simul. 7(3), 1192–1219 (2009).

20. Noé, F., Schütte, C., Vanden-Eijnden, E., Reich, L. & Weikl, T. R. Constructing the equilibrium ensemble of folding pathways from short off-equilibrium simulations. Proc. Natl. Acad. Sci. U.S.A. 106(45), 19011–19016 (2009).

21. Vanden-Eijnden, E. Transition-path theory and path-finding algorithms for the study of rare events. Annu. Rev. Phys. Chem. 61, 391–420 (2010).

22. Helfmann, L., Borrell, E. R., Schütte, C. & Koltai, P. Extending transition path theory: Periodically driven and finite-time dynamics. J. Nonlinear Sci. 30(6), 3321–3366 (2020).

23. Gupta, P. B. et al. Stochastic state transitions give rise to phenotypic equilibrium in populations of cancer cells. Cell 146(4), 633–644 (2011).

24. Geiß, C., Salas, E., Guevara-Coto, J., Régnier-Vigouroux, A. & Mora-Rodríguez, R. A. Multistability in macrophage activation pathways and metabolic implications. Cells 11(3), 404 (2022).

25. Castiglione, F., Tieri, P., Palma, A. & Jarrah, A. S. Statistical ensemble of gene regulatory networks of macrophage differentiation. BMC Bioinform. 17(Suppl 19), 506; 119–128 (2016).

26. Murray, P. J. Macrophage polarization. Annu. Rev. Physiol. 79, 541–566 (2017).

27. Eftimie, R. & Barelle, C. Mathematical investigation of innate immune responses to lung cancer: The role of macrophages with mixed phenotypes. J. Theor. Biol. 524, 110739; 110739 (2021).

28. Singhal, S. et al. Human tumor-associated monocytes/macrophages and their regulation of T cell responses in early-stage lung cancer. Sci. Transl. Med. 11(479), eaat1500 (2019).

29. Deligne, C. et al. Matrix-targeting immunotherapy controls tumor growth and spread by switching macrophage phenotype targeting microenvironmental cues to switch TAM phenotype. Cancer Immunol. Res. 8(3), 368–382 (2020).

30. Yang, Q. et al. The role of tumor-associated macrophages (TAMs) in tumor progression and relevant advance in targeted therapy. Acta Pharm. Sin. B 10(11), 2156–2170 (2020).

31. Malyshev, I. & Malyshev, Y. Current concept and update of the macrophage plasticity concept: Intracellular mechanisms of reprogramming and M3 macrophage “switch” phenotype. Biomed Res. Int. 2015, 341308; 2314–6133 (2015).

32. Kim, K. & Wang, J., Potential energy landscape and robustness of a gene regulatory network: Toggle switch. PLoS Comput. Biol. 3(3), e60 (2007). https://doi.org/10.1371/journal.pcbi.0030060

33. Yabo, Y. A., Niclou, S. P. & Golebiewska, A. Cancer cell heterogeneity and plasticity: A paradigm shift in glioblastoma. Neuro-oncology 24(5), 669–682 (2022).

34. Frank, A. S., Sikorski, A. & Röblitz, S. Spectral clustering of Markov chain transition matrices with complex eigenvalues. ArXiv: 2206.14537 (2022). Preprint available at https://doi.org/10.48550/arXiv.2206.14537.

