## Supplementary Material for "Macrophage phenotype transitions in a stochastic gene-regulatory network model"

#### Supplementary Method

##### *The deterministic ODE model*

The variable $x_1$ represents STAT1, a marker for M1 polarization, and the variable $x_2$ represents STAT6, a marker for M2 polarization, along with two input signals (denoted by $S_1$ and $S_2$). The equations of the ODE model read

$$x_1' = \big(a_1 \cdot H^+(x_1,k_1,n_1) + S_1\big) \cdot H^-(x_2,p_2,l_2) + b_1 - q_1 x_1, \tag{S.1}$$
$$x_2' = a_2 \cdot H^+(x_2,k_2,n_2) + S_2 \cdot H^-(x_1,p_1,l_1) + b_2 - q_2 x_2, \tag{S.2}$$

where $' = d/dt$. The term $H^+(x_i,k_i,n_i) = x^{n_i}/(x^{n_i} + {k_i}^{n_i})$ denotes a stimulatory Hill function, whereas $H^-(x_i,p_i,l_i) = p_i^{l_i}/(p_i^{l_i} + {x_i}^{l_i})$ denotes an inhibitory Hill function. The parameters $a_i, b_i, q_i \in \mathbb{R}$ are constants and defined in Table S1 and can be viewed in the network diagram in Fig. 2 of the manuscript. The equation for $x_2$ is based on the assumption that interferons inhibit activation of STAT6 by IL-4 in human monocytes by inducing SOCS-1 gene expression[1].

| Parameter | Description |
|---|---|
| $a_{1,2}$ | Strength of self-stimulation |
| $b_{1,2,}$ | Basal activation rates |
| $n_{1,2}$ | Exponents in the Hill functions for self-stimulation |
| $k_{1,2}$ | Thresholds in the Hill functions for self-stimulation |
| $l_{1,2}$ | Exponents in the Hill functions for mutual inhibition |
| $p_{1,2}$ | Thresholds in the Hill function for mutual inhibition |
| $q_{1,2}$ | Deactivation rates |
| $S_{1,2}$ | Input signal strength |

**Table S1.** Model Parameters in equations (S.1)–(S.2)

##### *Deriving the chemical master equation*

The chemical master equation (CME) is the most accurate description of a chemical reaction system if the number of reacting species is small. It takes into account random fluctuations in the time points and the order of reactions. For a system involving $D$ different species, the state space is $\mathbb{N}^D$, i.e. the space of $D$-dimensional positive integer vectors containing the copy-numbers of each of the $D$ species. The CME describes how a probability density in this space evolves over time. The CME can usually only be solved numerically, which is a tedious task. Alternatively, one can draw sample realizations of the underlying stochastic Markov jump process by using the stochastic simulation algorithm (SSA)[2].

In order to derive the CME from the deterministic ODE model, the ODE system is first decomposed into four elementary reactions:

$$\emptyset \xrightarrow{\alpha_1(x_1,x_2)} x_1, \quad \emptyset \xrightarrow{\alpha_2(x_1,x_2)} x_2, \quad x_1 \xrightarrow{\alpha_3(x_1)} \emptyset, \quad x_2 \xrightarrow{\alpha_4(x_2)} \emptyset,$$

with the four stochastic propensity functions

$$\begin{aligned}
\alpha_1(x_1;x_2) &= \left(a_1 \cdot H^+(x_1,k_1,n_1) + S_1\right) \cdot H^-(x_2,p_2,l_2) + b_1, \\
\alpha_2(x_1;x_2) &= a_2 \cdot H^+(x_2,k_2,n_2) + S_2 \cdot H^-(x_1,p_1,l_1) + b_2, \\
\alpha_3(x_1) &= q_1 \cdot x_1, \\
\alpha_4(x_2) &= q2 \cdot x_2,
\end{aligned}$$

and stoichiometric change matrix

$$S = \begin{pmatrix} 1 & 0 & -1 & 0 \\ 0 & 1 & 0 & -1 \end{pmatrix}.$$

Denote by $p(i,j)$ the probability to find the system in state $(x_1 = i, x_2 = j)$ at time $t$. The chemical master equation is the balance equation for the change in this probability density:

$$\begin{aligned}
\frac{d}{dt}p(i,j) = \quad & \alpha_1(i-1,j) \cdot p(i-1,j) + \alpha_3(i+1) \cdot p(i+1,j) \\
& + \alpha_2(i,j-1) \cdot p(i,j-1) + \alpha_4(j+1) \cdot (i,j+1) \\
& - [\alpha_1(i,j) + \alpha_3(i) + \alpha_2(i,j) + \alpha_4(j)] \cdot p(i,j)
\end{aligned}$$

#### *Molecular copy numbers*

To construct a stochastic model from the ODE system in equations (S.1)–(S.2), we need to know the volume of the system. A typical volume of a human alveolar macrophage is $4990 \pm 174\ \mu m^3$[3], which corresponds to a system volume of about $V = 5 \cdot 10^{-12}$L. Given that volume, molar concentrations in mol/L can be transformed into molecular copy numbers by multiplying them with $V \cdot n_A$, where $n_A$ denotes Avogadro's constant. This transformation applies to all initial values and to all parameters that contain units of concentration in mol/L.

When we constructed the ODE system (see equations (S.1)–(S.2)), $x_1$ and $x_2$ were assumed to be dimensionless. Hence, before we could apply the transformation into copy numbers, we needed to re-introduce the dimension mol/L by multiplying the initial values and all parameters that contain units of $x_1$ or $x_2$, i.e. $a_1, a_2, b_1, b_1, k_1, k_2, p_1, p_2, S_1, S_2$, with a typical unit value, which we have chosen to be $u = 0.43 \cdot 10^{-11}$ mol/L ($= 4.3$ nmol/mL)[1]. The definition of each parameter is provided in the Table in S1.

#### *Latin hypercube sampling*

To explore the parameter space and resulting dynamics, we employ Latin Hypercube Sampling (LHS)[4,5] to generate samples of parameter values. LHS differs from random sampling in that it simultaneously stratifies on all input dimensions[6]. Because we wish to sample a broad range of parameters, we apply LHS to the exponents (base 10) to cover a broad range of parameter values. In our work, this translates to dividing each of the parameter exponent ranges into equi-probable intervals and choosing one sample from each defined hypercube. Because we have no a priori knowledge of the parameter values, we assume a uniform distribution.

We sample the exponents of the parameters $a_1$, $a_2$, $b_1$, $b_2$, $k_1$, $k_2$, $q_1$, $q_2$, $s_1$, $s_2$ over orders of magnitude ranging from $10^{-3}$ to $10^3$. These parameters were selected because in our previous work[7], they were shown to be sensitive using a variance-based sensitivity analysis. For each sample of parameter values, we record the number of eigenvalues of $Q$ near 0 (we define a threshold of less than .001). This tells us the number of potential clusters for that parameter set. Because this is a necessary but not sufficient condition for a corresponding number of clusters, we then apply PCCA+ for those sets which have two or more small eigenvalues. We record the parameter sets which correspond to two or more clusters. For 80,000 parameter samples, we find only bistable parameter sets.

#### *Transition path theory*

TPT takes as input the transition rate matrix $Q = (\ell_{ij})_{i,j \in \mathscr{S}} = Q(i,j)_{i,j \in \mathscr{S}}$ of a Markov jump process $\{X\}_{t \in \mathbb{R}}$ defined on a discrete state space $\mathscr{S}$. Based on the assumption that the process $\{X\}_{t \in \mathbb{R}}$ is irreducible and ergodic, the time-reversed process $\{\tilde{X}\}_{t \in \mathbb{R}}$ is also a Markov jump process with a rate matrix $\tilde{Q} = (\tilde{\ell})_{i,j \in \mathscr{S}}$ given by $\tilde{\ell}_{i,j} = \frac{\pi_j}{\pi_i} \ell_{ji}$[8], where $\pi$ is the invariant distribution (or stationary density) of both processes $\{X\}_{t \in \mathbb{R}}$ and $\{\tilde{X}\}_{t \in \mathbb{R}}$.

TPT is based on *reactive trajectories*, which are defined as transitions between two non-empty, disjoint subsets $A$ and $B$ of the state-space $\mathscr{S}$, where one set is the source and the other one is the sink state[8,9].

The discrete forward $q^+$ and backward $q^-$ committor functions are essential to calculate statistical properties and dynamical information of the reactive trajectories. The forward committor $q^+$ is defined by[8], as follows,

[1] If experimental measurement values are available, the value of $u$ can be fitted to those values.

$$\begin{cases} \sum_{j\in\mathscr{S}} \ell_{ij} q_j^+ = 0 & \forall i \in (A\cup B)^c \\ q_j^+ = 0 & \forall i \in A, \\ q_j^+ = 1 & \forall i \in B, \end{cases} \tag{S.3}$$

and the backward commitor $q^-$ as

$$\begin{cases} \sum_{j\in\mathscr{S}} \tilde{\ell}_{ij} q_j^- = 0 & \forall i \in (A\cup B)^c \\ q_j^- = 1 & \forall i \in A, \\ q_j^- = 0 & \forall i \in B. \end{cases} \tag{S.4}$$

Given the above definitions of committor functions, rate matrices and stationary densities, we can calculate statistical properties that allow for a dynamical interpretation of the ensemble of reactive trajectories between subsets $A$ and $B$ in the state space $\mathscr{S}$[10]. Table S2 summarizes important measures in TPT to describe dynamical information and states the corresponding interpretation.

For any $i \in \mathscr{S}$, the *distribution of reactive trajectories* $\mu^R = (\mu_i^R)_{i\in\mathscr{S}}$ is given by (see[8])

$$\mu_i^R = \pi_i q_i^+ q_i^- . \tag{S.5}$$

Normalizing $\mu_i^R$ by *the probability to be on a transition at time t* (i.e., $Z_{AB} = \sum_{j\in\mathscr{S}} \pi_j q_j^+ q_j^-$)[10], one defines the *normalized distribution of reactive trajectories* ($\mu_i^{AB}$)[8], as

$$\mu_i^{AB} = Z_{AB}^{-1}\, \mu_i^R. \tag{S.6}$$

Two other important statistics are the *net average number of reactive trajectories observed per time unit* ($f_{ij}^+$) and the *average number of transitions from A to B per time unit* ($k_{AB}$)[8].

The definitions of $f_{ij}^+$ and $k_{AB}$ build on the *probability current of reactive trajectories*, $f_{ij}^{AB}$, which is defined in[8] as follows: For all pairs of states $(i,j), i,j \in \mathscr{S}, i \neq j$,

$$f_{ij}^{AB} = \pi_i q_i^- \ell_{ij} q_j^+, \quad \text{where } \ell_{ij} = Q(i,j). \tag{S.7}$$

$f_{ij}^{AB}$ describes the average rate at which a reaction trajectory transitions from state $i$ to state $j$. Moreover, the *transition rate* from $A$ to $B$, $k_{AB}$, is given by

$$k^{AB} = \underbrace{\sum_{i\in A, j\in\mathscr{S}} f_{ij}^{AB}}_{k^{A\to}} = \underbrace{\sum_{i\in\mathscr{S}, j\in B} f_{ij}^{AB}}_{k^{\to B}}, \tag{S.8}$$

where $k^{A\to}$ and $k^{\to B}$ are the discrete rates of leaving $A$ and entering $B$, respectively (see[8,10]). Equality in equation (S.8) holds for stationary Markov processes $\{X_t\}_{t\in\mathbb{R}}$[10]. The *effective current* $f_{ij}^+$ is defined by (see[8])

$$f_{ij}^+ = \max(f_{ij}^{AB} - f_{ji}^{AB}, 0). \tag{S.9}$$

Furthermore, the *average time duration of a transition from A to B*, $t^{AB}$, is defined in[10] by

$$t^{AB} := \frac{Z^{AB}}{k^{AB}}. \tag{S.10}$$

While the above characteristics are defined on the entire state space $\mathscr{S}$, we consider for the following definitions the coarse-grained sets from the Markov State Model, which is defined by matrix $P_c(\tau)$. For the coarse partition of the state space $\tilde{\mathscr{S}} = \{S_1, ..., S_n\}$, where $A$ and $B$ are identical to individual $S_i$ or a combination of several $S_i$, *the coarse-grained flux from set $S_i$ to set $S_f$*, is defined, according to[9], by

$$\tilde{F}_{S_i \to S_f} = \sum_{k\in S_i, l\in S_f} f_{k,l}^{AB} \tag{S.11}$$

Given the coarse-grained flux $\tilde{F}_{S_i\to S_f}$, we can now define *individual pathways* $P_i$ connecting A and B[9], by connecting neighbouring sets $S_i, i \in [n]$, with $S_1 = A$ and $S_n := B$, such that $\tilde{F}_{S_i\to S_f} \neq 0$.

Given the set of all individual pathways $P_i$ from $A$ to $B$, we describes the *relative probability* of a path segment from $S_i$ to $S_f$ (similar to[9]) by

$$p^r_{S_i \to S_f} = \frac{\tilde{F}_{S_i \to S_f}}{\sum\limits_{i=S_i, f \neq S_i} \tilde{F}_{S_i \to S_f}}, \tag{S.12}$$

where the sum represent the total flow from state $S_i$ to neighboring states $S_j$, $j \in [n]$, $j \neq i$ on the $A \to B$ transition paths.

Finally, the relative probabilities of the path segments $p^r_{S_i \to S_f}$ along an individual pathway $P_i$ from $A$ to $B$ are combined to define the *total relative probability* $p^r_i$ *of path* $P_i$.

Alternatively, there exist several, so called, *Path Decomposition Algorithms*[8,9] that allow the decomposition into individual pathways $P_i$. However, if used on the entire state space $\mathscr{S}$ they can be computationally very expensive. Due to our relative small coarse-grained state space $\tilde{\mathscr{S}}$, we decided to calculate the decomposition manually, as described above.

### Supplementary Tables

| Symbol | Name | Interpretation |
|---|---|---|
| Entire state space | | |
| $A \to B$ | Reactive trajectory from $A$ to $B$ | A trajectory that leaves $A$ and enters $B$ without returning to $A$ in between |
| $q^+$ | Forward committor | Prob. that a trajectory initiated in A will reach B before A |
| $q^-$ | Backward committor | Prob. that a trajectory initiated in B will reach A before B |
| $f_{ij}^{AB}$ | Prob. current of reactive trajectories | Average rate at which a reactive trajectory from $A$ to $B$ transitions from state $i$ to state $j$ |
| $f_{AB}^{+}$ | Effective current | $f_{ij}^{+} = \max(f_{ij}^{AB} - f_{ji}^{AB}, 0)$ |
| $\mu_i^{AB}$ | Prob. distribution of reactive trajectories | Prob. that a reactive trajectory passes through state $i$ |
| $t_{AB}$ | Transition time | Average time duration of a transition from $A$ to $B$ |
| $k^{A\to}$ | Transition rate out of $A$ | Average rate at which trajectories leave $A$ |
| $k^{\to B}$ | Transition rate into $B$ | Average rate at which trajectories enter $B$ |
| CG state space | | |
| $\tilde{\pi}_S$ | Total stationary prob. of macrostate $S$ | |
| $\tilde{F}_{S_i \to S_f}$ | Coarse-grained flux between macrostates $S_i$ and $S_f$ | |
| $\tilde{T}_{S_i \to S_f}$ | Transition probability between macrostates $S_i$ and $S_f$ | Inter-macrostate transitions (based on source-state $S_i$) |

**Table S2. Nomenclature of Transition Path Theory (TPT)**. Table presents the dynamical information about the ensemble of reactive trajectories and their interpretations. Details on TPT can be found in the Methods section and the Supplementary Methods online.

| Case | Num. of states | Eigenvalues of $Q_c$ | Cluster weights | Transition state matrix | Transition prob. matrix |
|---|---|---|---|---|---|
| 1 | 2 | $\lambda_1 = 0$<br>$\lambda_2 = -0.9884$<br>$(\lambda_3 = -0.9997)$ | $\omega_1 = 0.10031$<br>$\omega_2 = 0.89969$ | $Q_c = 10^{-4} \cdot \begin{pmatrix} 0.4707 & -0.4707 \\ -0.6109 & 0.6109 \end{pmatrix}$ | $P_c = \begin{pmatrix} 0.9981 & 0.0019 \\ 0.0000 & 1.0000 \end{pmatrix}$ |
| 2 | 2 | $\lambda_1 = 1.000$<br>$\lambda_2 = 0.9962$<br>$(\lambda_3 = 0.0008)$ | $\omega_1 = 0.099103$<br>$\omega_2 = 0.9009$ | $Q_c = 10^{-4} \cdot \begin{pmatrix} -0.3860 & 0.3860 \\ 0.0625 & -0.0625 \end{pmatrix}$ | $P_c = \begin{pmatrix} 0.9962 & 0.0038 \\ 0.0000 & 1.0000 \end{pmatrix}$ |
| 3 | 2 | $\lambda_1 = 0$<br>$\lambda_2 = -0.0214$<br>$(\lambda_3 = -4.6872)$ | $\omega_1 = 0.45748$<br>$\omega_2 = 0.54252$ | $Q_c = \begin{pmatrix} -0.0269 & 0.0269 \\ 0.0272 & -0.0272 \end{pmatrix}$ | $P_c = \begin{pmatrix} 0.1243 & 0.8757 \\ 0.0068 & 0.9932 \end{pmatrix}$ |
| 4 | 2 | $\lambda_1 = 0$<br>$\lambda_2 = -0.0021$<br>$(\lambda_3 = -0.9261)$ | $\omega_1 = 0.097708$<br>$\omega_2 = 0.90229$ | $Q_c = \begin{pmatrix} -0.0024 & 0.0024 \\ 0.0001 & -0.0001 \end{pmatrix}$ | $P_c = \begin{pmatrix} 0.8094 & 0.1906 \\ 0.0001 & 0.9999 \end{pmatrix}$ |
| 5 | 3 | $\lambda_1 = 0,$<br>$\lambda_2 = -2.8653 \cdot 10^{-5}$<br>$\lambda_3 = -6.1945 \cdot 10^{-4}$<br>$(\lambda_4 = -0.9902)$ | $\omega_1 = 0.016006$<br>$\omega_2 = 0.084884$<br>$\omega_3 = 0.89911$ | $Q_c = 10^{-3} \cdot \begin{pmatrix} -0.0327 & 0.0033 & 0.0294 \\ 0.6020 & -0.6075 & 0.0055 \\ -0.0042 & 0.0043 & -0.0001 \end{pmatrix}$ | $P_c = \cdot \begin{pmatrix} 0.9957 & 0.0014 & 0.0029 \\ 0.0580 & 0.9414 & 0.0006 \\ 0.0000 & 0.0000 & 1.0000 \end{pmatrix}$ |

**Table S3. Clustering results from PCCA+.** Table shows the number of metastable states (i.e., phenotypes), the calculated eigenvalues ($\lambda_i, i = \{1,2,3,4\}$) and gaps, cluster weights ($\omega_i,\ i = \{1,2\}$), transition rate matrix $Q_c$, and transition probability matrix $P_c$ for the respective five parameter sets ("Case").

| Case 1: Forward | | Case 1: Backward | | Case 2: Forward | | Case 2: Backward | |
|---|---|---|---|---|---|---|---|
| Start: | H/L | Start: | H/H | Start: | L/H | Start: | H/L |
| End: | H/H | End: | H/L | End: | H/L | End: | L/H |
| **Statistics: Average time duration per transition** | | | | | | | |
| $t$ | 3.508329e-01 | $t$ | 1.888526e-01 | $t$ | 1.745843e-01 | $t$ | 5.180507e-01 |
| **Coarse-grained flux directions and amount:** | | | | | | | |
| $\mathbf{S_i \rightarrow S_f}$ | $\mathbf{\tilde{F}_{S_i \rightarrow S_f}}$ | $\mathbf{S_i \rightarrow S_f}$ | $\mathbf{\tilde{F}_{S_i \rightarrow S_f}}$ | $\mathbf{S_i \rightarrow S_f}$ | $\mathbf{\tilde{F}_{S_i \rightarrow S_f}}$ | $\mathbf{S_i \rightarrow S_f}$ | $\mathbf{\tilde{F}_{S_i \rightarrow S_f}}$ |
| $H/L \rightarrow H/H$ | 0 | $H/H \rightarrow H/L$ | 0 | $L/H \rightarrow H/L$ | 0 | $H/L \rightarrow L/H$ | 0 |
| $H/L \rightarrow C$ | 4.030848e-14 | $H/H \rightarrow C$ | 7.307069e-14 | $L/H \rightarrow C$ | 2.126413e-14 | $H/L \rightarrow C$ | 1.236879e-14 |
| $H/H \rightarrow H/L$ | 0 | $H/L \rightarrow H/H$ | 0 | $H/L \rightarrow L/H$ | 0 | $L/H \rightarrow H/L$ | 0 |
| $H/H \rightarrow C$ | 0 | $H/L \rightarrow C$ | 0 | $H/L \rightarrow C$ | 0 | $L/H \rightarrow C$ | 0 |
| $C \rightarrow H/L$ | 0 | $C \rightarrow H/H$ | 0 | $C \rightarrow L/H$ | 0 | $C \rightarrow H/L$ | 0 |
| $C \rightarrow H/H$ | 1.404820e-14 | $C \rightarrow H/L$ | 2.257171e-14 | $C \rightarrow H/L$ | 7.004957e-15 | $C \rightarrow L/H$ | 3.946240e-15 |
| **Transition paths and transition probabilities** | | | | | | | |
| $H/L \rightarrow H/H$ | 0% | $H/H \rightarrow H/L$ | 0% | $L/H \rightarrow H/L$ | 0% | $H/L \rightarrow L/H$ | 0% |
| $H/L \rightarrow C \rightarrow H/H$ | 100% | $H/H \rightarrow C \rightarrow H/L$ | 100% | $L/H \rightarrow C \rightarrow H/L$ | 100%. | $H/L \rightarrow C \rightarrow L/H$ | 100% |
| Case 3: Forward | | Case 3: Backward | | Case 4: Forward | | Case 4: Backward | |
| Start: | L/L | Start: | H/L | Start: | L/L | Start: | L/H |
| End: | H/L | End: | L/L | End: | L/H | End: | L/L |
| **Statistics: Average time duration per transition** | | | | | | | |
| $t$ | 1.776616e-07 | $t$ | 1.395691e-06 | $t$ | 4.367060e-01 | $t$ | 1.784335e-01 |
| **Coarse-grained flux directions and amount:** | | | | | | | |
| $\mathbf{S_i \rightarrow S_f}$ | $\mathbf{\tilde{F}_{S_i \rightarrow S_f}}$ | $\mathbf{S_i \rightarrow S_f}$ | $\mathbf{\tilde{F}_{S_i \rightarrow S_f}}$ | $\mathbf{S_i \rightarrow S_f}$ | $\mathbf{\tilde{F}_{S_i \rightarrow S_f}}$ | $\mathbf{S_i \rightarrow S_f}$ | $\mathbf{\tilde{F}_{S_i \rightarrow S_f}}$ |
| $L/L \rightarrow H/L$ | 1.232997e-04 | $L/L \rightarrow H/L$ | 0 | $L/L \rightarrow L/H$ | 0 | $L/L \rightarrow L/H$ | 0 |
| $L/L \rightarrow C$ | 5.619505e-09 | $L/L \rightarrow C$ | 0 | $L/L \rightarrow C$ | 1.009425e-14 | $L/L \rightarrow C$ | 0 |
| $H/L \rightarrow L/L$ | 0 | $H/L \rightarrow L/L$ | 9.039354e-05 | $L/H \rightarrow L/L$ | 0 | $L/H \rightarrow L/L$ | 0 |
| $H/L \rightarrow C$ | 0 | $H/L \rightarrow C$ | 2.660313-08 | $L/H \rightarrow C$ | 0 | $L/H \rightarrow C$ | 2.052825e-14 |
| $C \rightarrow L/L$ | 0 | $C \rightarrow L/L$ | 4.933935e-10 | $C \rightarrow L/L$ | 0 | $C \rightarrow L/L$ | 6.876109e-15 |
| $C \rightarrow H/L$ | 7.050022e-11 | $C \rightarrow H/L$ | 0 | $C \rightarrow L/H$ | 3.554568e-15 | $C \rightarrow L/H$ | 0 |
| **Transition paths and transition probabilities:** | | | | | | | |
| $L/L \rightarrow H/L$ | 99.99% | $H/L \rightarrow L/L$ | 99.97% | $L/L \rightarrow L/H$ | 0% | $L/H \rightarrow LL$ | 0% |
| $L/L \rightarrow C \rightarrow H/L$ | 0.01% | $H/L \rightarrow C \rightarrow L/L$ | .03% | $L/L \rightarrow C \rightarrow L/H$ | 100% | $L/H \rightarrow C \rightarrow LL$ | 100% |

**Table S4. Transition Path Theory (TPT) results for sets of two phenotype states (two each of L/L, H/L, L/H, H/H).** Each subcase specifies the start and end sets of the transition directions, as well as complement (C) set, as well as the average time duration ($t$) it takes for a phenotype to switch. Based on the coarse-grained transition fluxes between metastable (phenotype) sets (e.g., $S_i$ indicates the initial set and $S_f$ the final set), all possible transition paths and their probabilities are calculated as described in the Methods section. See Supplementary Table S2 online for description and interpretation of TPT nomenclature. Methodological details are presented in the Methods section and in Supplementary Methods online.

| Subcase 1 | | Subcase 2 | | Subcase 3 | |
|---|---|---|---|---|---|
| Start: | L/L | Start: | L/L & H/L | Start: | L/L |
| End: | L/H & H/L | End: | L/H | End: | L/H |
| Compl.: | C | Compl.: | C | Compl.: | C |
| Inter.: | - | Inter.: | - | Inter.: | H/L |
| **Statistics: Average time duration per transition** | | | | | |
| $t$ | 1.645091e−01 | $t$ | 4.010152e−01 | $t$ | 2.031077e+00 |
| **Coarse-grained flux directions and amount:** | | | | | |
| $\mathbf{S_i \to S_f}$ | $\tilde{\mathbf{F}}_{\mathbf{S_i \to S_f}}$ | $\mathbf{S_i \to S_f}$ | $\tilde{\mathbf{F}}_{\mathbf{S_i \to S_f}}$ | $\mathbf{S_i \to S_f}$ | $\tilde{\mathbf{F}}_{\mathbf{S_i \to S_f}}$ |
| $L/L \to H/L$ | 4.283462e−14 | $L/L \to C$ | 3.6797e−14 | $L/L \to H/L$ | 6.947925e−17 |
| $L/L \to C$ | 3.765562e−14 | $H/L \to C$ | 3.913517e−15 | $L/L \to C$ | 3.679917e−14 |
| $C \to H/L$ | 7.501706e$^{-}$16 | $C \to L/H$ | 1.136061e−14 | $H/L \to H/L$ | 2.360166e−11 |
| $C \to L/H$ | 1.049604e−14 | | | $H/L \to C$ | 1.576060e−16 |
| | | | | $C \to L/H$ | 1.051028e−14 |
| | | | | $C \to H/L$ | 5.286661e−17 |
| **Transition paths and transition probabilities** | | | | | |
| $L/L \to C \to L/H$ | 43.66% | $L/L \to C \to L/H$ | 90.39% | $L/L \to C \to L/H$ | 99.31% |
| $L/L \to H/L$ | 53.22% | $H/L \to C \to L/H$ | 9.61% | $L/L \to H/L \to C \to L/H$ | 1.2615e-6% |
| $L/L \to C \to H/L$ | 3.12% | | | $L/L \to C \to H/L \to C \to L/H$ | 3.3135e-6% |
| Subcase 4 | | Subcase 5 | | Subcase 6 | |
| Start: | L/L | Start: | L/H | Start: | H/L |
| End: | H/L | End: | H/L | End: | L/H |
| Compl.: | C | Compl.: | C | Compl.: | C |
| Inter.: | L/H | Inter.: | L/L | Inter.: | L/L |
| **Statistics: Average time duration per transition** | | | | | |
| $t$ | 1.353744e+05 | $t$ | 6.990304e+03 | $t$ | 1.359403e+04 |
| **Coarse-grained flux directions and amount:** | | | | | |
| $\mathbf{S_i \to S_f}$ | $\tilde{\mathbf{F}}_{\mathbf{S_i \to S_f}}$ | $\mathbf{S_i \to S_f}$ | $\tilde{\mathbf{F}}_{\mathbf{S_i \to S_f}}$ | $\mathbf{S_i \to S_f}$ | $\tilde{\mathbf{F}}_{\mathbf{S_i \to S_f}}$ |
| $L/L \to H/L$ | 4.292451e−14 | $L/H \to C$ | 2.185682e−14 | $H/L \to L/L$ | 5.811830e−15 |
| $L/L \to C$ | 8.647682e−16 | $L/L \to H/L$ | 8.438635e−15 | $H/L \to C$ | 5.821236e−15 |
| $L/H \to C$ | 3.139963e−14 | $L/L \to C$ | 1.250078e−14 | $L/L \to C$ | 2.650196e−14 |
| $L/H \to L/H$ | 8.209541e−07 | $L/L \to L/L$ | 2.208563e−09 | $L/L \to L/L$ | 2.285861e−09 |
| $C \to H/L$ | 7.515452e−16 | $C \to H/L$ | 1.421303e−17 | $C \to L/H$ | 3.960937e−15 |
| $C \to L/H$ | 8.916429e−16 | $C \to L/L$ | 1.310601e−14 | $C \to L/L$ | 8.115767e−15 |
| **Transition paths and transition probabilities:** | | | | | |
| $L/L \to H/L$ | 98.30% | $L/H \to C \to H/L$ | 0.11% | $H/L \to C \to L/H$ | 16.41% |
| $L/L \to C \to H/L$ | 0.90% | $L/H \to C \to L/L \to C \to H/L$ | 6.2193e−7% | $H/L \to L/L \to C \to L/H$ | 1.8999e−4% |
| $L/L \to C \to L/H \to C \to H/L$ | $1.87e-8$% | $L/H \to C \to L/L \to H/L$ | 3.8166e−4% | $H/L \to C \to L/L \to C \to L/H$ | 1.2783e−4% |

**Table S5. Transition Path Theory (TPT) results for three phenotype states (L/L, H/L and L/H).** Each subcase specifies the start and end sets of the transition directions, as well as intermittent (I) or complement (C) set, as well as the average time duration $(t)$ it takes for a phenotype to switch. Based on the coarse-grained transition fluxes between metastable (phenotype) sets (e.g., $S_i$ indicates the initial set and $S_f$ the final set), all possible transition paths and their probabilities are calculated as described in the Methods section. See Supplementary Table S2 online for description and interpretation of TPT nomenclature. Methodological details are presented in the Methods section and in Supplementary Methods online. Abbreviations: "Compl." refers to the complement set (C); "Inter." stands for intermittent set (I).

### Supplementary Figures

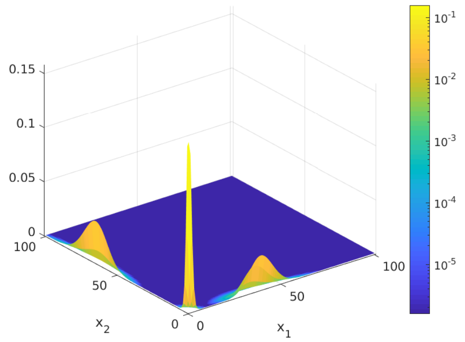


**Figure S1. Three macrophage phenotypes**: Surface plot of the sum of the three partial densities corresponding to the three identified metastable clusters. The ratio between $x_1, x_2$ characterize the phenotype: Low activation in $x_1$ and high activation level in $x_2$ describes the Low/High (L/H) phenotype. Low activation in $x_1$ and low activation level in $x_2$ describes the Low/Low (L/L) phenotype, and high activation in $x_1$ and low activation level in $x_2$ describes the High/Low (H/L) phenotype.

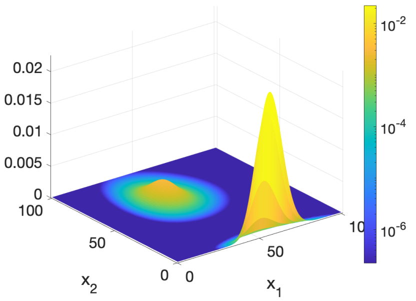


**(a)** Case 1: H/H and H/L phenotype

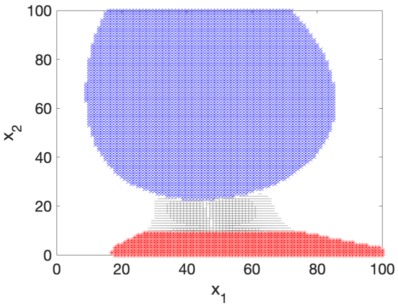


**(b)** Case 1: Transitions (grey) from H/L (red) to H/H (blue)

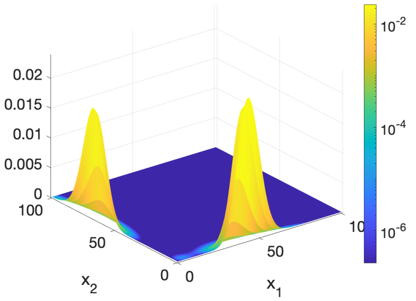


**(c)** Case 2: H/L and L/H phenotype

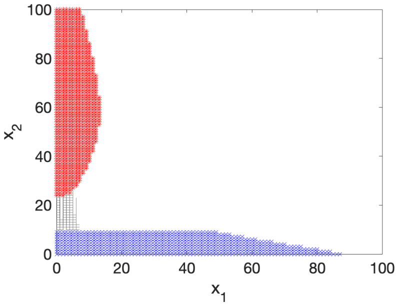


**(d)** Case 2: Transitions (grey) from L/H (red) to H/L (blue)

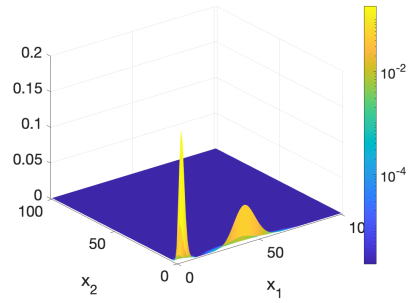


**(e)** Case 3: L/L and H/L phenotype

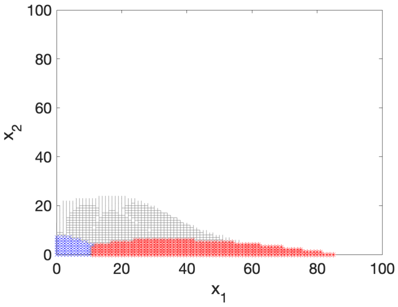


**(f)** Case 3: Transitions (grey) from H/L (red) to L/L (blue)

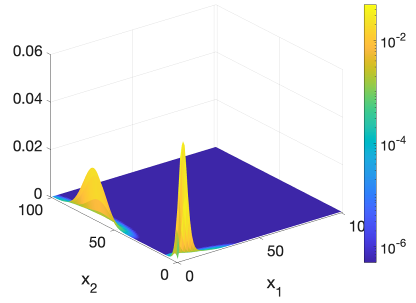


**(g)** Case 4, Partial Densities

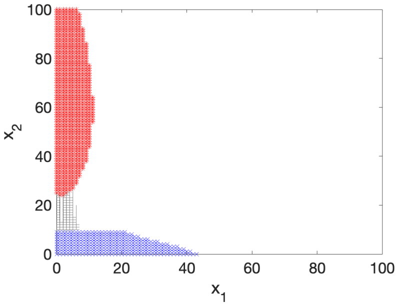


**(h)** Case 4: Transitions (grey) from L/H (red) to L/L (blue)

**Figure S2. Bistable phenotype configurations.** The left panel shows a surface plot of the sum of the partial densities which correspond to two metastable clusters. The ratio between $x_1, x_2$ defines the phenotypes. In (a) we see high activation levels in both $x_1$ and $x_2$, and thus can classify a High/High (H/H) phenotype. High activation in $x_1$ and low activation in $x_2$, describes a H/L phenotype. Similarly, we see bistable clusters of phenotypes in (c) H/L and L/L, in (e) L/L and H/L, and in (g) L/L and L/H. The right panel shows calculated transition path flow directions in (b), (d), (f) and (h). In these transition flow graphs, the red colored set indicates the starting set and the blue colored set the ending set. The gray colored area represents the transition flow.

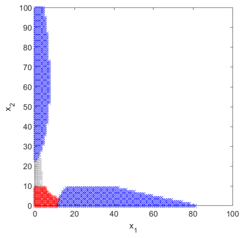

**(a)** Subcase 1: Transition flow paths (grey) from L/L (red) to H/L (blue) and L/H (blue)

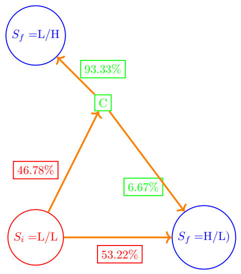


**(b)** Subcase 1: Probabilities between L/L (red) and H/L (blue) and L/H (blue) phenotypes

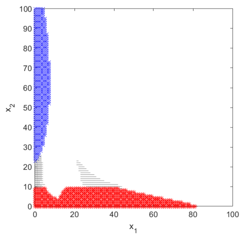

**(c)** Subcase 2: Transition (grey) from L/L (red) and H/L (red) to L/H (blue)

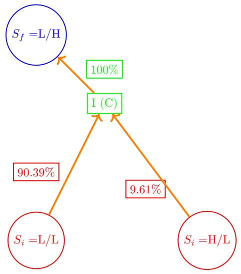


**(d)** Subcase 2: Probabilities between L/L (red) and H/L (red) and L/H (blue) phenotypes

**Figure S3. Transition flows, paths and probabilities between three phenotypes with combined start and end sets.** The left column shows the transition path flow (in grey color) in the CME state-space between a given start set (in red color) and end set (in blue color). The right column represents the transition path flows in a transition network together with the corresponding path segment probabilities, which are added next to the arrows of the flow direction. Red colored circles represent the start sets ($S_i$, with i for initial), and blue ones the end sets ($S_f$, with f for final). Green color refers to the state space without any phenotype, i.e., the complement set (C) to the states ($S_i$ and $S_f$). The path segment probabilities were calculated from equation (4) in the Methods section. Their color refers to the set from where the transition flow comes. Supplementary Table S5 online gives a quantitative overview over the transition dynamics, including all possible transition paths and their respective probabilities. Subcases 3–6 are presented in Fig. 4 in the article.

### References


1. Dickensheets, H. L., Venkataraman, C., Schindler, U. & Donnelly, R. P. Interferons inhibit activation of STAT6 by interleukin 4 in human monocytes by inducing SOCS-1 gene expression. Proc. Natl. Acad. Sci. U.S.A. **96**(19), 10800–10805 (1999).
2. Gillespie, D. T. A general method for numerically simulating the stochastic time evolution of coupled chemical reactions.. J. Comput. Phys. **22**(4), 403–434 (1976).
3. Kromback, F. et al. Cell size of alveolar macrophages: An interspecies comparison.. Environ. Heal. Perspect. **105**(Suppl 5), 1261–1263 (1997).
4. Stein, M. Large sample properties of simulations using Latin hypercube sampling.. Technometrics **29**(2), 143–151 (1987).
5. McKay, M. D.,Beckman, R. J. & Conover, W. J. A comparison of three methods for selecting values of input variables in the analysis of output from a computer code. Technometrics **42**(1), 55–61 (2000).
6. Loh, W. On Latin hypercube sampling. Ann Stat. **24**(5), 2058–2080 (1996).
7. Frank, A. S., Larripa, K., Ryu, H., Snodgrass, R. G. & Röblitz, S. Bifurcation and sensitivity analysis reveal key drivers of multistability in a model of macrophage polarization. J. Theor. Biol. **509**, 110511; 1–15 (2021).
8. Metzner, P., Schütte, C. & Vanden-Eijnden, E. Transition path theory for Markov jump processes. Multiscale Model. Simul. **7**(3), 1192–1219 (2009).
9. Noé, F., Schütte, C., Vanden-Eijnden, E., Reich, L. & Weikl, T. R. Constructing the equilibrium ensemble of folding pathways from short off-equilibrium simulations. Proc. Natl. Acad. Sci. U.S.A. **106**(45), 19011–19016 (2009).
10. Helfmann, L., Borrell, E. R., Schütte, C. & Koltai, P. Extending transition path theory: Periodically driven and finite-time dynamics. J. Nonlinear Sci. **30**(6), 3321–3366 (2020).
